# Combinatorial perturbation sequencing on single cells using microwell-based droplet random pairing

**DOI:** 10.1101/2022.08.03.502265

**Authors:** Run Xie, Yang Liu, Shiyu Wang, Xuyang Shi, Zhantao Zhao, Longqi Liu, Ya Liu, Zida Li

**Author notes:** Contributed equally (Run Xie, Yang Liu).

## Abstract

Combinatorial drug therapy reduces drug resistance and disease relapse, but informed drug combinations are lacking due to the high scale of possible combinations and the relatively simple phenotyping strategies. Here we report combinatorial perturbation sequencing (CP-seq) on single cells using microwell-base droplet random pairing. CP-seq uses oligonucleotides to barcode drugs, encapsulates drugs and cells in separate droplets, and pairs cell droplets with two drug droplets randomly on a microwell array chip to complete combinatorial drug treatment and barcode-tagging on cells. The subsequent single-cell RNA sequencing simultaneously detects the single-cell transcriptomes and drug barcodes to demultiplex the corresponding drug treatment. The microfluidic droplet operations had robust performance, with overall success rate among the microwells being up to 83%. We then progressively validated the CP-seq by performing single drug treatment and then combinatorial drug treatment. Leveraging the advantage of droplet microfluidics in massive multiplexing, the CP-seq can test thousands of drug combinations in a single experiment and represents a great technology for combinatorial perturbation screening with high throughput and comprehensive profiling.

## Introduction

Current drug discovery primarily aims to find agents that target a specific signaling pathway. However, due to the redundancy of cell signaling and the inter-cellular heterogeneity, using these drugs alone sometimes leads to suboptimal results. Relapse may arise due to previously unaware drug resistance existing in the cell population.^1^

Combinatorial drug therapy using multiple targets can potentially serve as a better treatment strategy.^2^ By simultaneously targeting multiple mechanisms, combinatorial drug therapy can significantly reduce the chance of relapse.^3^ Indeed, the efficacy of this strategy has been demonstrated in the application of treating bacterial infection by using multiple antibiotics jointly.^4^ In addition, the efficacy of combinatorial drug treatment has been shown in cancer treatment, where a few combinatorial chemotherapies have been approved by the Food and Drug Administration of the United States.^5^

Nevertheless, drug combinations with informed efficacy are still limited, primarily due to the formidable workload in performing combinatorial screening. For example, the number of two-drug combinations of a library with 1,000 drugs are almost half a million, and the number of combinations of three or more drugs can be even more huge, making it costly both workload-wise and finance-wise. In addition, currently the characterization of drug screening experiments mainly relies on a few simple phenotypes, such as proliferation, morphology, and a handful of biomarkers, barely offering a comprehensive response profile. Finally, current drug screening experiments are mostly performed on bulk samples, providing the response on population average. Given the ubiquitous cellular heterogeneity, cells respond to drugs differently even among the same type, and the existence of a rare subtype may result in drug resistance and disease relapse. Therefore, analysis on single-cell level is desirable.^6^

A few works have been reported to address these technical challenges in drug screening. Microfluidics has been adopted to develop high throughput combinatorial screening using programmable microvalves,^7-10^ microwells,^11^ micropillars,^12^ or multilayered channels.^13^ However, these methods used simple phenotypes such as proliferation rate as the readouts. To address that, transcriptome analysis has been proposed to characterize cell responses to drug perturbations in a much more comprehensive manner,^14^ and single-cell RNA sequencing (scRNA-seq) has been proposed as a means to further tease out the heterogeneous response of among the cell populations, provided that the perturbation information can be properly indexed into the sequencing results. To this end, barcoding strategies based on oligo transfection,^15^ single-nucleotide polymorphism,^16^ and nuclear hashing^17^ have been developed to examine the transcriptomic response to perturbations on single-cell level. Nevertheless, the operations of these techniques relied on microtiter plates as the reactors. Given the scale of combinatorial perturbation screening, both the workload and reagent consumption can be unbearable when used for such applications. Therefore, techniques that can perform single-cell transcriptomic analysis on combinatorial perturbations are yet to be developed.

Leveraging the advantage of droplet microfluidics in developing high throughput assays,^11, 18^ here we report combinatorial perturbation sequencing (CP-seq) which can simultaneously perturb cells with thousands of drug combinations and perform single-cell transcriptomic analysis. CP-seq encapsulates cells and oligo-coded drugs in microdroplets, respectively, and performs droplet pairing in specially designed microwells, where two drug droplets randomly pair with one cell droplet, to complete the combinatorial perturbation, before the cells are subject to scRNA-seq. Since the drug barcodes are compatible with scRNA-seq workflow, the perturbation of individual cells can be recovered from the sequencing data. We first demonstrated the robust performance of the droplet operation, showing that droplet pairing had a high overall success rate among the microwells of 83%. We then validated CP-seq by first performing treatment of single drugs with known effects, showing the effectiveness of the experimental process. We further validated CP-seq by imposing combinatorial drug treatment, and the scRNA-seq results confirmed the efficacy. CP-seq is able to perform drug treatment of thousands of combinations on a glass-slide-sized microfluidic chip, and the capability can be further scaled up by making larger chips. We envision that CP-seq would serve as a versatile tool for high throughput screening with comprehensive profiling on single-cell level and greatly facilitate drug discovery.

## Methods

### Design and fabrication of microfluidic devices

Three microfluidic devices were used, including two droplet generators for cell and drug encapsulation, respectively, and one microwell array device for droplet pairing. Droplet generators adopted flow-focusing structures driven by a negative pressure, and the channel heights was 90 and 40 μm throughout for cell and drug encapsulation, respectively. The microwell array device was composed of a layer with microwell arrays and a layer with flow chamber. The depths of the microwells and the flow chamber were 70 μm and 220 μm, respectively, in the device for combinatorial treatment. In the device for single drug treatment, the depths of large and small microwell were 80 and 50 μm, respectively. Other dimensions are shown in **Figure S1**.

All microfluidic layers except the flow chamber were fabricated through replica molding of polydimethylsiloxane (PDMS) using SU-8 molds. Fabrication of the SU-8 molds used photolithography and was outsourced to a microfabrication company (HICOMP MicroTech, Suzhou). The SU-8 molds were then silanized in vacuum chamber overnight, before PDMS prepolymer (10:1 base-to-hardener ratio) was poured on the mold, degassed, and baked in a 90 °C oven for 2 h. The cured PDMS was then peeled off and cut into desired size, before reservoir, inlets, and outlets were punched. The diameters of oil reservoirs, aqueous phase reservoirs, and inlets/outlets were 10, 5, and 1.5 mm, respectively. PDMS surfaces were then treated with oxygen plasma (PDC-002, Harrick Plasma), placed in contact, and baked in 110 °C for 30 minutes to complete bonding. The mold for the flow chamber layer of the microwell array device used a slightly different but more economical method. Double-sided tape (3M; 140 μm in thickness) was laminated on polyethylene terephthalate membrane (80 μm in thickness), and the two layers were laser-cut and sticked to a Petri dish on the tape side, which was then used as the mold.

### Cell culture and preparation

Human breast cancer cell line (MCF-7) was used for the cell study. Cells were cultured in Dulbecco’s modified Eagle’s medium (DMEM) supplemented with 10% fetal bovine serum (FBS) and 1% penicillin-streptomycin (PS) in a humidified incubator (IP610, Yamato Scientific) with 5% CO2 and 100% humidity at 37 °C. Cell were passaged at 80% confluency. Cell suspension was prepared by spinning down cells, washing with DPBS, and resuspending in a working buffer (DMEM with 6% Ficoll-PM400, 5% FBS, 2.5% HEPES, 1% PS, and 1% F68). Cells counting and viability test were performed using AO/PI staining on a fluorescence cell analyzer (Shanghai Ruiyu Biotech., China), and cell density of 8000 cells/μL was adopted for the generation of cell droplets.

### Reagent preparation

Reagent information is listed in **Table S1**. Four drugs were used in the experiment, namely doxorubicin hydrochloride (DOX), fluorouracil (5-FU), cyclophosphamide (CP), and paclitaxel (PTX). DOX and CP were dissolved in water of molecular biology-grade, and 5-FU and PTX were dissolved in dimethyl sulfoxide (DMSO) following supplier’s suggestion. Three different concentrations of each drug were used, and the concentrations are listed in **Table S2**.

### Design and preparation of the concanavalin A-based drug barcode (CADB) complex

The drug barcode consists of a PCR handle, a unique 10–base barcode, and a 21-base poly(A) tail. Full sequences are listed in **Table S3**. The oligonucleotides were synthesized and biotinylated by BGI Tech Solutions (Beijing). Upon arrival, the biotinylated oligonucleotides were diluted in nuclease-free water at the concentration of 100 nM. Biotinylated concanavalin A and streptavidin were dissolved in 50% glycerol at concentration of 1.6 μM separately. To assemble the CADB complex, streptavidin was firstly mixed with biotinylated oligonucleotides and incubated for 10 minutes at room temperature, before biotinylated concanavalin A was added and incubated for 10 min at room temperature.

### Droplet generation and manipulation

Droplets were generated by filling reservoirs with either oil or the aqueous phase and applying negative pressure at the outlet. Negative pressure was applied by pulling an air-filled syringe. The 87 μm cell droplets, 43 μm drug droplets, and 50 μm drug droplets were generated by pulling a 30 mL plastic syringe (302833, BD) from the initial position of 15 mL to 20 mL, 15 mL to 20 mL, and 17 mL to 20 mL respectively. A cryovial (431386, Corning) was punched and connected between the syringe and the device outlet for droplet collection. The generated cell droplets were stored on ice, and the generated drug droplets were gently pooled for later use. After loading into the microwell array device, droplets were merged by treating the PDMS device with corona for 5 seconds using a handheld corona treater (BD-20AC, Electro-Technic Products, Chicago). Droplets were then retrieved into a centrifuge tube for incubation. After incubation, droplets were demulsified by slowly adding 1H,1H,2H,2H-perfluoro-1-octanol (PFO) into the oil. After washing, cells were resuspended in PBS solution containing 0.04% BSA.

### Single-cell RNA sequencing (scRNA-seq)

The DNBelab C4 (MGI Tech Co., Shenzhen) was used for single-cell RNA sequencing library preparation, as previously described.^19^ Key steps included cell and bead encapsulation, emulsion breakage, bead collection, reverse transcription, and cDNA amplification to generate barcoded libraries. The sequencing libraries were quantified using the Qubit ssDNA Assay Kit (Q10212, Invitrogen), and sequencing was performed on a high throughput sequencer (DIPSEQ T1, MGI Tech Co., Shenzhen).

### Analysis of scRNA-seq data

Raw read data in FASTQ format were first processed using in-house Perl scripts to obtain correspondence between cells and drug treatment conditions. The FASTQ data of single-cell transcriptome were aligned to the genome of human reference genome (GRCh38) through STAR^20^, and the unique molecular identifier (UMI) count matrix was generated using PISA (v0.12a; https://github.com/shiquan/PISA). The gene expression matrix was processed using Seurat (v4.1.0).^21^ Cells with mitochondrial gene counts greater than 20% and cells expressing fewer than 1000 or more than 10,000 genes were excluded. The filtered data were then normalized and scaled using default parameters. The top 2000 highly variable genes for each library were used for further processing. All datasets were then integrated using the “merge” function in Seurat. Finally, we conducted dimension reduction for the scaled merged dataset by PCA analysis. The first 30 principal components were used to construct a K-nearest-neighbor graph using the “FindNeighbors” function, and the cell clusters were assigned through the “FindClusters” function. Visualization was shown using UMAP. Up- and down-regulated genes for each time point cluster cells versus all combination cells were computed using the “FindMarkers” function in Seurat package, and marker genes with the adjusted p-value < 0.05, average log2FC > 1.5, and average log2FC < −1.5 were screened for further analysis. Volcano plots were then used to display the result of the differential gene expression. Gene ontology analysis, particularly for cellular component ontology, was applied to explore the biological significances of combinatorial drug treated cells using “clusterProfiler” package (v4.2.2).^22^

## Results

### Workflow of the CP-seq technology

To achieve multiplexity, CP-seq uses oligonucleotide (oligo) sequences to code the drug information and uses concanavalin A (ConA), which is a glycoprotein-binding protein applied in cell membrane labeling^23^, as the linker to tag the oligo barcodes to cell membranes, as shown in **Figure 1a**. The oligo mainly consists of a PCR handle, a unique 10-base barcode to store drug information, and a 21-base poly(A) tail for downstream capturing. By conjugating ConA with oligo, the complex of ConA-based drug barcode (CADB) is created. Each drug solution is mixed with a designated CADB complex that encodes the drug condition, including the drug molecule and dose, and the mix is encapsulated into droplets, as shown in **Figure 1b**. All drug droplets are then pooled, resulting in a drug library in the form of droplets. Meanwhile, cells are encapsulated into droplets that are larger than the drug droplets. Since CP-seq does not require single-cell encapsulation, an additional step of droplet sorting is not necessary.

**Figure 1.**
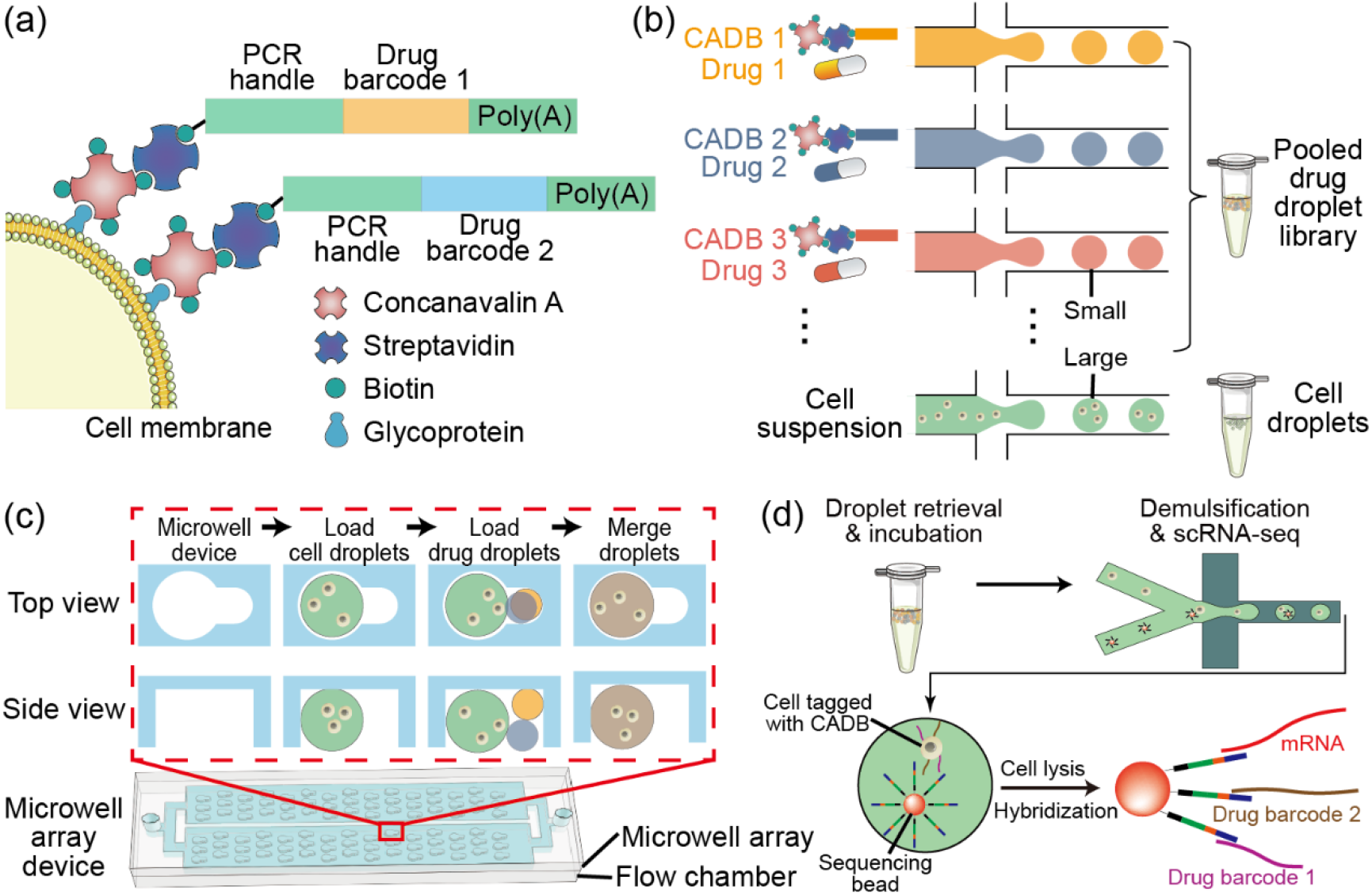
Overview of the combinatorial perturbation sequencing technology. (a) Schematic showing the principle of cell tagging with the drug barcode. Concanavalin A, which is conjugated with a designated oligonucleotide sequence, binds to the cell membrane and thus tags the cell. (b) Barcoded drug droplets and cell droplets with differential diameters are generated using microfluidics. Drugs are pre-mixed with the concanavalin A-based drug barcode (CADB). (c) Cell droplets are loaded into the microwell array device and occupy the large microwells, before drug droplets are loaded and occupy the small microwells. Each small microwell randomly captures two drug droplets. Droplets in each microwell unit are merged via corona treatment, resulting in combinatorial drug treatment and CADB tagging on cells. (d) Droplets are then incubated and demulsified to collect cells for single-cell RNA sequencing (scRNA-seq). The sequencing bead simultaneously capture mRNA and drug barcodes through poly(T) tail.

The combinatorial drug treatment is performed in a microfluidic device consisting of an upper layer of microwell array facing down and a lower layer of flow chamber (**Figure 1c**; **Figure S1**; **Video S1**). Each microwell unit consists of two interconnected microwells, with a diameter of 90 and 50 μm, respectively. The depth of the microwells is 70 μm. A glass slide-sized chip houses roughly 26800 microwell units. Cell droplets, with diameters of 87 μm, are first loaded into the flow chamber. Since water is much lighter than the continuous phase of oil (∼1.6 g/mL), the droplets float into the microwell and occupy the large half of each microwell unit, leaving out the small half. Drug droplets, with smaller diameters of 43 μm, are then loaded into the flow chamber and float into the small half of each microwell unit. Since the microwells are relatively deep, with a proper quality control on the droplet dimension, each microwell captures precisely two drug droplets. The cell droplet and the two drug droplets are merged using corona treatment. The droplets are then retrieved and incubated to complete the combinatorial drug treatment, before the cells are subject to single-cell RNA sequencing (scRNA-seq). Sequencing beads capture both the mRNA and drug barcode oligos by the functionalized poly(T) on the bead surface, as shown in **Figure 1d**, and consequently, both the transcriptome and the two drug barcodes can be recovered from the sequencing data.

### Performance of the microfluidic manipulation of droplets

Robust performance of droplet manipulation is the foundation of the CP-seq technology. Therefore, we first sought to characterize and optimize the operation parameters related to droplet manipulation. Uniform droplets were generated using flow-focusing devices driven by a negative pressure at the outlet, and the resultant cell and drug droplets were 86.8 ± 3.7 and 42.3 ± 1.5 μm in diameters, respectively, with variations less than 5% (**Figure 2a&b**; **Figure S1**). Using a cell density of 8000 cells/μL, 97.9% of the cell droplets contained one or more cells, with 67% containing two to four cells.

**Figure 2.**
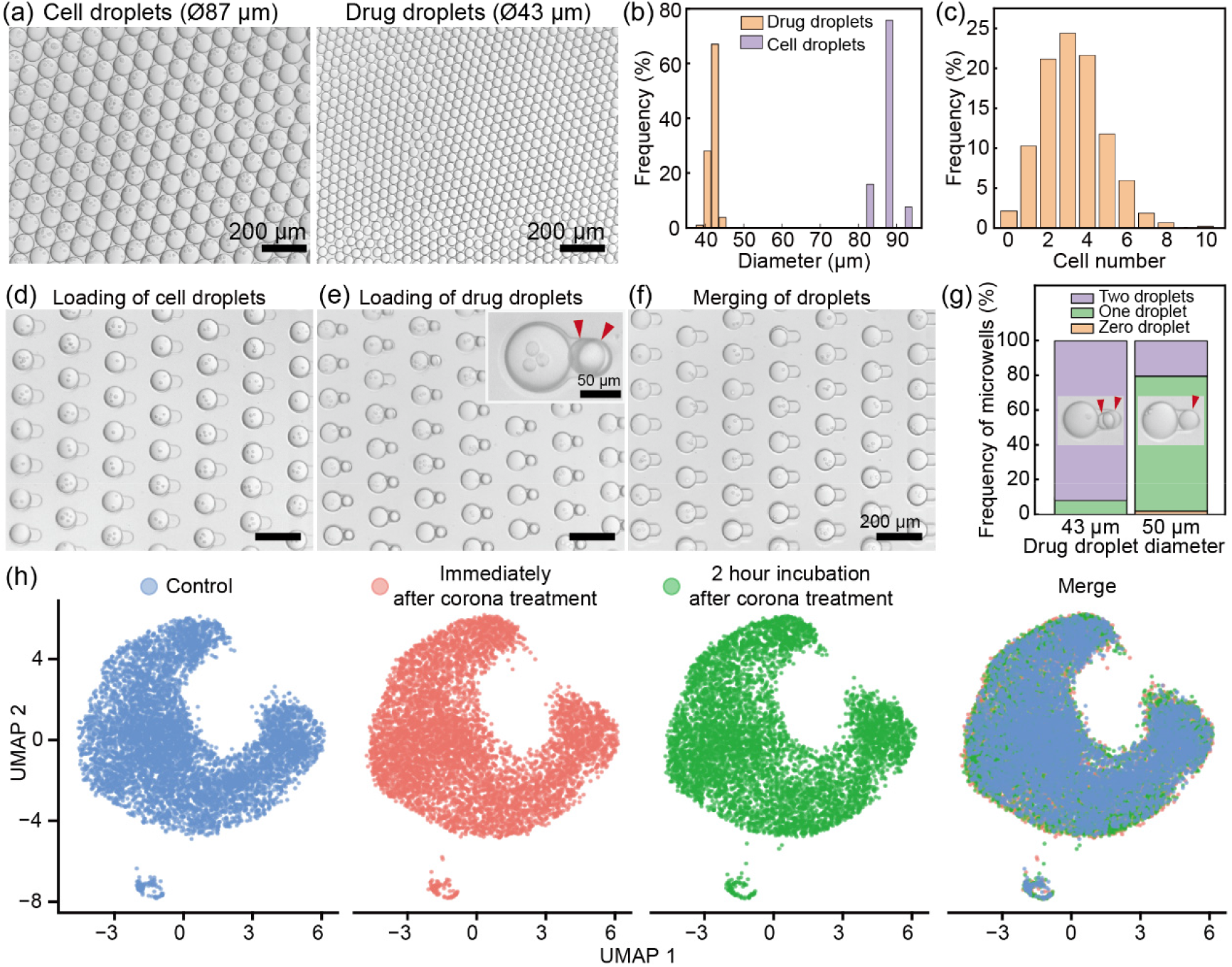
Performance of the droplet manipulation platform. (a) Micrographs of the generated cell droplets and drug droplets. (b) Diameter distribution of the cell droplets and drug droplets. (c) Distribution of the cell numbers in each cell droplet. (d-f) Micrographs showing the process of cell droplet loading, drug droplet loading, and droplet merging, with microwell utilization rates being 99%, 92%, and 90%, respectively. Simultaneous capturing of two drug droplets can be identified under bright-field microscope, as indicated by red arrows. (g) Frequency of microwells occupied by different numbers of droplets when using droplets of different diameters. (h) UMAP plot of the scRNA-seq data showing the effect of the corona treatment on cells.

The droplet trapping and pairing on the microwell array device also showed robust performance. After cell droplet loading, more than 99% of the microwells were occupied by a cell droplet (**Figure 2d**). After drug droplet loading, more than 92% of the microwells captured two drug droplets (**Figure 2e**). The microwell array device was then subject to a brief corona treatment, resulting in interface destabilization and subsequent droplet merging. By slightly tilting the device and ensuring that all three droplets were in contact, more than 90% of the microwells were merged after the corona treatment (**Figure 2f**). By flipping over the device, nearly all the droplets floated out of the microwells and could be collected at the outlet (**Video S1**). In the process of droplet pairing, the size of the drug droplets was an important parameter. For example, when drug droplets had a diameter of 50 μm instead of 43 μm, only 20.4% microwells captured two droplets, and 77.7% microwells captured one droplet (**Figure 2g**).

Though cell culture in droplets has been well studied,^24^ corona treatment is a less standard operation. Corona treatment generates an instant but strong electric potential, which may impose disturbance on the gene expression of the cells. Therefore, we specifically investigated the effect on cells of the dose of corona treatment in our experiment setting, which was 5 seconds. We performed scRNA-seq on MCF-7 cells immediately and two hours after 5 seconds’ corona treatment and compared them with those not treated with corona at all. After dimensionality reduction using Uniform Manifold Approximation and Projection (UMAP), the transcriptome data were highly overlapping among the three groups, suggesting that the brief treatment with corona had negligible effect on cells (**Figure 2h**).

### Validation of the CP-seq using single drug treatment

We then sought to validate CP-seq’s performance in profiling single cell transcriptomes on drug perturbations. The gene expression after the treatment of a specific chemotherapy drug has been well studied, and it offers a good reference for comparison. Therefore, we first performed single drug treatment using the CP-seq technology to validate the processes involving scRNA-seq and drug demultiplexing.

The microwell array device in CP-seq was adapted by reducing the depth of the small microwell to allow the trapping of single drug droplet (**Figure 3a**). Consequently, more than 97% of the microwell units captured a cell droplet and a single drug droplet (**Figure 3b**). Using this microwell array device, we first characterized the performance CADB binding and recovering using qPCR. We designed oligo with poly(A) tail, the accompanying reverse transcription primer, and PCR amplification primers (**Table S4**). The oligo was conjugated to ConA, and the resultant CADB complex was used as the “drug” to treat and tag cells following the CP-seq experimental protocol. After “drug” treatment, instead of performing scRNA-seq, reverse transcription and qPCR was performed on the cell lysis to quantify the oligo. The oligo was successfully detected, and the qPCR results showed that on average about 18600 oligo molecules were recovered from each cell, consistent with reported works (**Figure S2**).^25^ These results suggested that the droplet manipulation was fully operational and the CADB-based cell-labelling and barcoding was successful.

**Figure 3.**
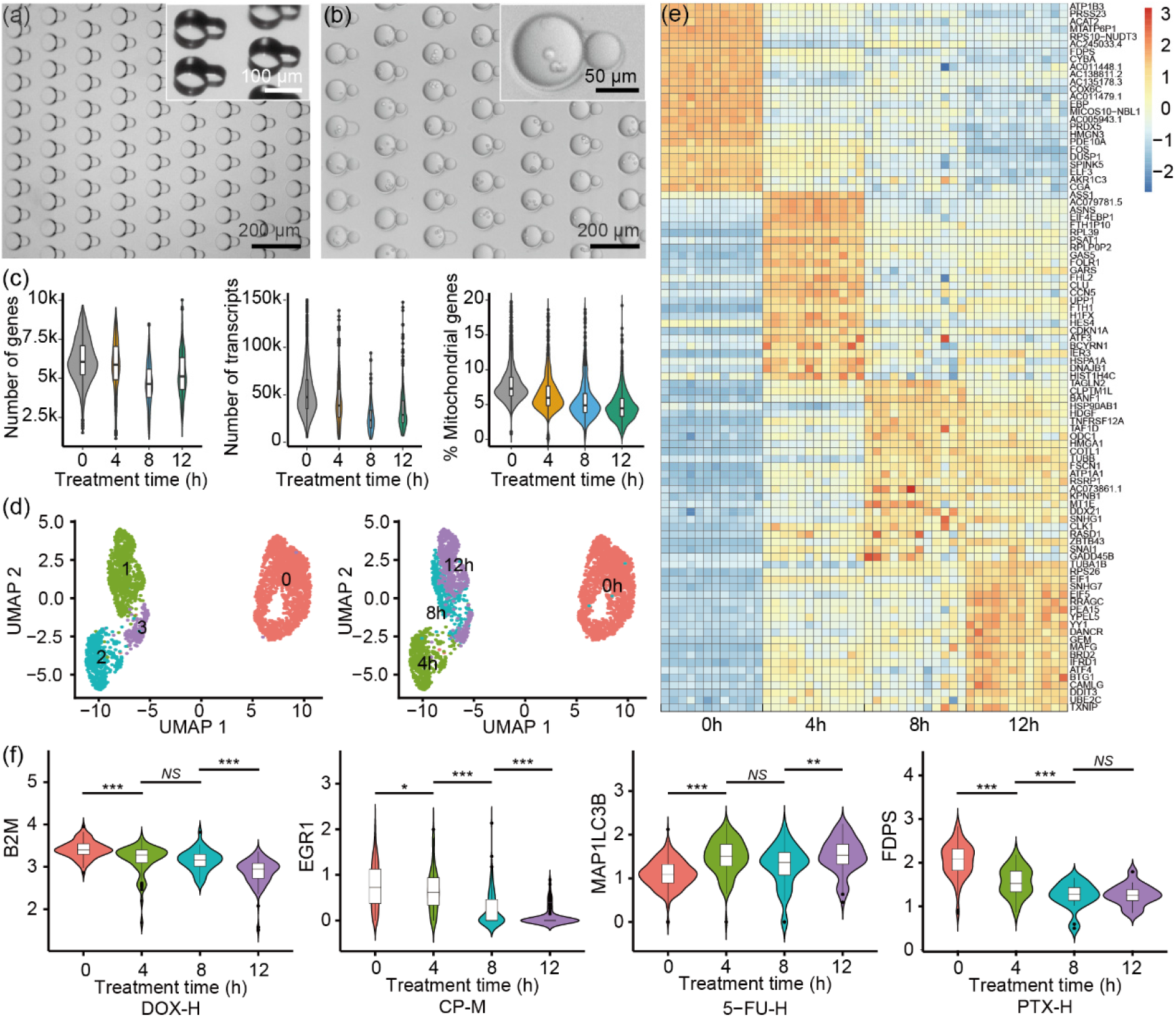
Validation of the CP-seq using single drug treatment. (a) Micrograph showing the microwell device used to capture single drug droplet. Microwells with differential depths were designed. Inset: close-up view of the microwell unit under a stereomicroscope. (b) Micrograph showing the capturing of cell droplets and single drug droplets. (c) Quality assessment of the scRNA-seq results under drug treatment of 0, 4, 8, and 12 hours. (d) Scatter plot of UMAP analysis showing the cell clustering. (e) Heatmap showing the gene expression at different time point. Each column represents a drug treatment condition at corresponding time point. (f) Temporal gene expression of representative genes. *, p < 0.05. **, p < 0.01. ***, p < 0.001. *N*.*S*., not significant.

We then sought to perform single drug treatment on cells. We chose breast cancer cell (MCF-7) and four commonly used chemotherapy drugs, namely doxorubicin (DOX), paclitaxel (PTX), 5-flurouracil (5-FU), and cyclophosphamide (CP), for the study. Three doses for each drug, suffixed with L (low), M (medium), and H (high), were incorporated in the experiments, resulting in 13 groups including the vehicle control (**Table S2**). In addition, we sought to profile the single-cell transcriptome with different incubation time to obtain the temporal development of gene expression. To obtain the suitable timeframe of drug treatment, we first performed cell viability assay since live cells were necessary for scRNA-seq. The results showed the viability had a sharp drop upon 16 hours’ treatment (**Figure S3**). Therefore, we chose 4, 8, and 12 hours as the incubation times.

We performed the CP-seq with the 13 treatment groups, and each drug condition was designated with a CADB barcode shown in **Table S3**. After drug treatment, the cells were incubated for 0, 4, 8, and 12 hours as separate experiments, and the cells then went through scRNA-seq. The scRNA-seq results showed good quality. The numbers of genes detected in each cell on average ranged from 4000 to 6000, the numbers of transcripts were more than 15000, and the percentages of mitochondrial genes were around 5% in experiments of different incubation times (**Figure 3c**). After alignment and error correction, we detected 3243 cells with corresponding drug treatment recovered. We then performed dimensionality reduction using uniform manifold approximation and projection (UMAP) to visualize the transcriptomes of cells incubated for different times. As shown in **Figure 3d**, the drug-treated cells showed distinct transcriptomes compared to the control group (0 h). Among drug-treated cells, those of the same incubation time were near each other, and the clustering results reasonably agreed with the experiment conditions of incubation time. We further visualized the temporal development of the average expression of the top 100 genes using a heatmap (**Figure 3e**). The heatmap showed the gradual change over time of the gene expression profiles. We then examined a few genes with known consequences upon drug treatments, and the results agreed with the established drug knowledge. For example, as shown in **Figure 3f**, the down-regulation of the B2M and EGR1 gene due to DOX and CP treatment were both observed^26^, and the up-regulation of the MAP1LC3B gene, which was responsive to DNA damage, due to 5-FU treatment was also observed.^27^ We also observed gene regulations that were previously not reported, such as the down-regulation of the FDPS gene upon PTX treatment, which might hold biological insights and could be further investigated.

### Validation of the CP-seq using combinatorial drug treatment

We then performed the complete process of the CP-seq for further validation. We used the above-mentioned 12 drug conditions (four drugs by three different doses) and a blank control for the experiments, and the incubation time was shortened to one hour, given the low viability of cells on combinatorial drug treatment. We additionally profiled the transcriptomes of cells immediately after drug treatment but without incubation for comparison.

We first sought to examine the drug barcoding performance. The number of combinations of two among the 13 drug conditions is 78; therefore, we expected to recover 78 barcode combinations from the sequencing data. In the no-incubation group, we recovered 71 combinations, as visualized in the chord diagram in **Figure 4a**. Seven combinations were missing, likely because of the data filtering in postprocessing which discarded transcriptomes of low quality. In the incubation group, we recovered 47 treatment combinations, which were fewer than that in the no-incubation group, presumably because of the cell death induced by the combinatorial treatment. We then compared the gene expression patterns in these two experiment groups using Gene Ontology (GO) enrichment analysis. Since the four drugs used in the experiments were all related to DNA damage and cell apoptosis and had similar effect on cells, we pooled the experiment groups for the analysis. In particular, we adopted cellular component ontology (CCO) terms to analyze the differences in cellular components between these two experiment groups. The results showed that the GO-CCO terms enriched in the group without incubation were mainly related to chromosomes in the nucleus, the genes of which perform functions such as mitosis (**Figure 4b**). In contrast, the GO terms enriched in the group with 1-hour incubation were mainly ribosomal and mitochondrial components, likely indicating a high cell stress. We then sought to examine the transcriptomic impact of combinatorial drug treatment compared to single drug treatment. For the ease of analysis, we pooled the data of each experiment groups and analyzed the differential gene expression. As shown in **Figure 4c**, the analysis visualized the up-regulation of tens of genes and the down-regulation of about 500 genes, which demonstrated the capability of the CP-seq in profiling gene expression under combinatorial drug treatment.

**Figure 4.**
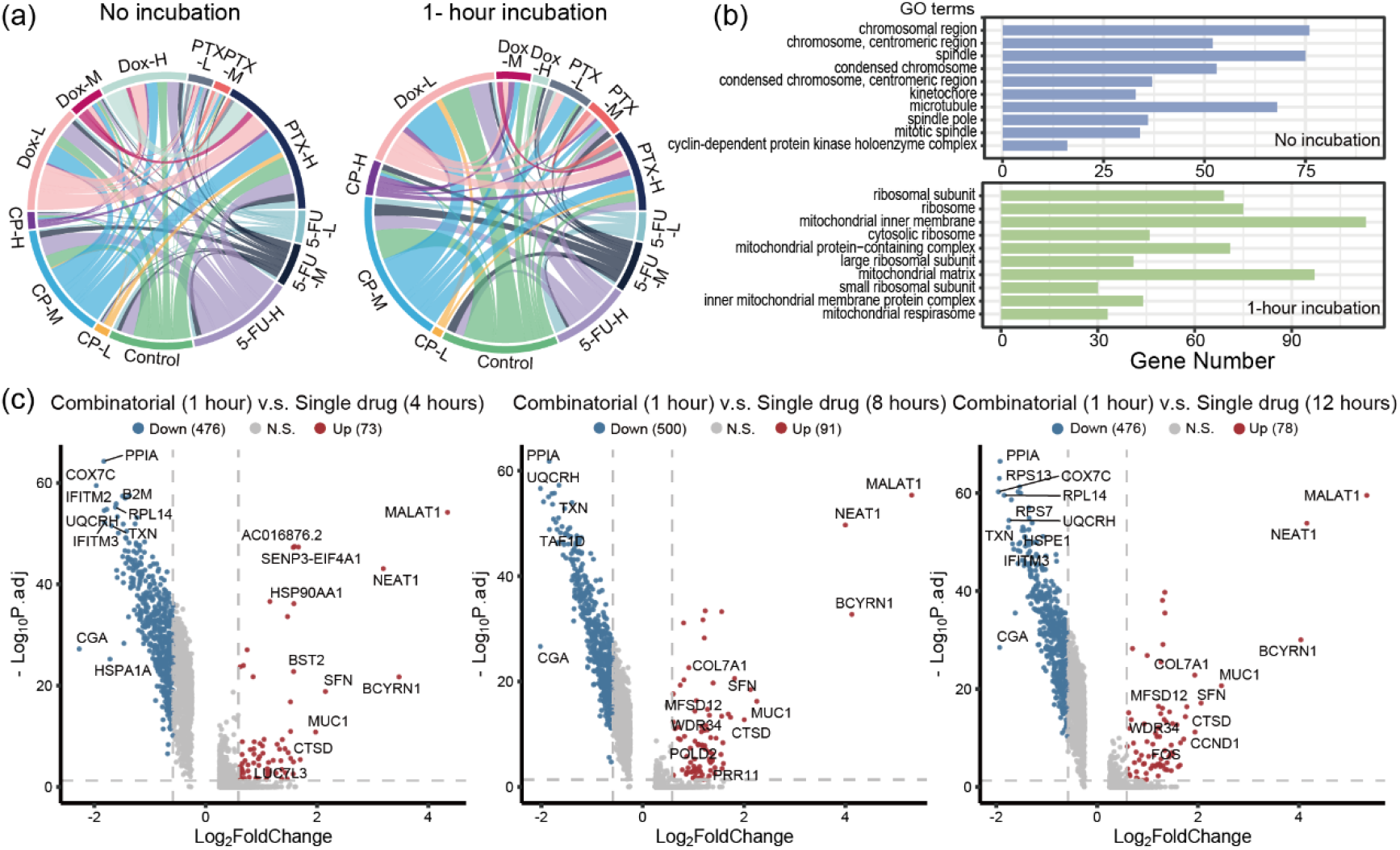
Validation of the CP-seq using combinatorial drug treatment. (a) Chord diagrams showing the drug combinations recovered from the sequencing data in groups with no incubation and 1-hour incubation. (b) Histogram showing the top ten Gene Ontology (GO) terms of cell component assigned to the above two transcriptomes. Results were ranked based on the adjusted p-value. (c) Volcano plots showing the differential gene expression of cells with combinatorial drug treatment for an hour compared to single drug treatment for 4 hours, 8 hours, and 12 hours. Genes that have an adjusted p-value smaller than 0.05 and an absolute value of fold change larger than 1.5 were considered significant.

## Conclusions and discussion

In this work, we developed a technology, named combinatorial perturbation sequencing (CP-seq), to obtain single-cell transcriptomes upon combinatorial drug treatments using droplet microfluidics. Using oligonucleotides for drug barcoding and a microwell array device for random combinatorial drug treatment, CP-seq can collect the transcriptomes of single cells treated with thousands of drug combinations in a single experiment. We optimized the droplet manipulation processes and validated the CP-seq by performing single drug treatment. We also performed combinatorial drugs treatments of 13 drug conditions and recovered 71 drug combinations out of the 78 possible combinations, showing the potential of CP-seq in finding new efficacious drug combinations. In addition to combinations of two drugs, CP-seq holds the versatility of being adapted to test combinations of multiple drugs by incorporating additional small microwells in each microwell unit.

Nevertheless, though the throughput of CP-seq can be easily scaled up by designing larger devices of microwell arrays, the generation of the drug droplet library can be labor-intensive if the drug library contains thousands of drugs, since the droplets of each drug are currently generated separately. A strategy to resolve this challenge is to adopt techniques that generate droplets with high automation. For example, the commercial instruments of microarray spotters can be utilized to generate high-volume droplet library, as demonstrated in **Figure S4** and by reported works.^28^ With such droplet generation techniques, drug droplet library of high volume can be prepared with minimal human intervention.

## Data availability

Sequencing data of this study are available in the CNGB Nucleotide Sequence Archive (https://db.cngb.org/cnsa/; accession number CNP0003242).

### Author contributions

R.X., Yan.L., S.W., L.L., YaL., and Z.L. conceptualized the project and designed the experiments. R.X., Yan.L., and Z.Z. performed the experiments. R.X., S.W., and X.S. analyzed the data. R.X., YaL., and Z.L. wrote the manuscript. All authors contributed to the manuscript.

## Acknowledgments

The authors thank Xiaoxiang Hu and Prof. Yujuan Chai for sharing equipment. This work was supported by the National Key Research and Development Program of China (2021YFF1200500), Guangdong Basic and Applied Basic Research Foundation (2021A1515110459), Shenzhen Overseas Talent Program, and Shenzhen University Faculty Startup Grant.

