## Supplementary Document for "Combinatorial perturbation sequencing on single cells using microwell-based droplet random pairing"

[#]Contributed equally

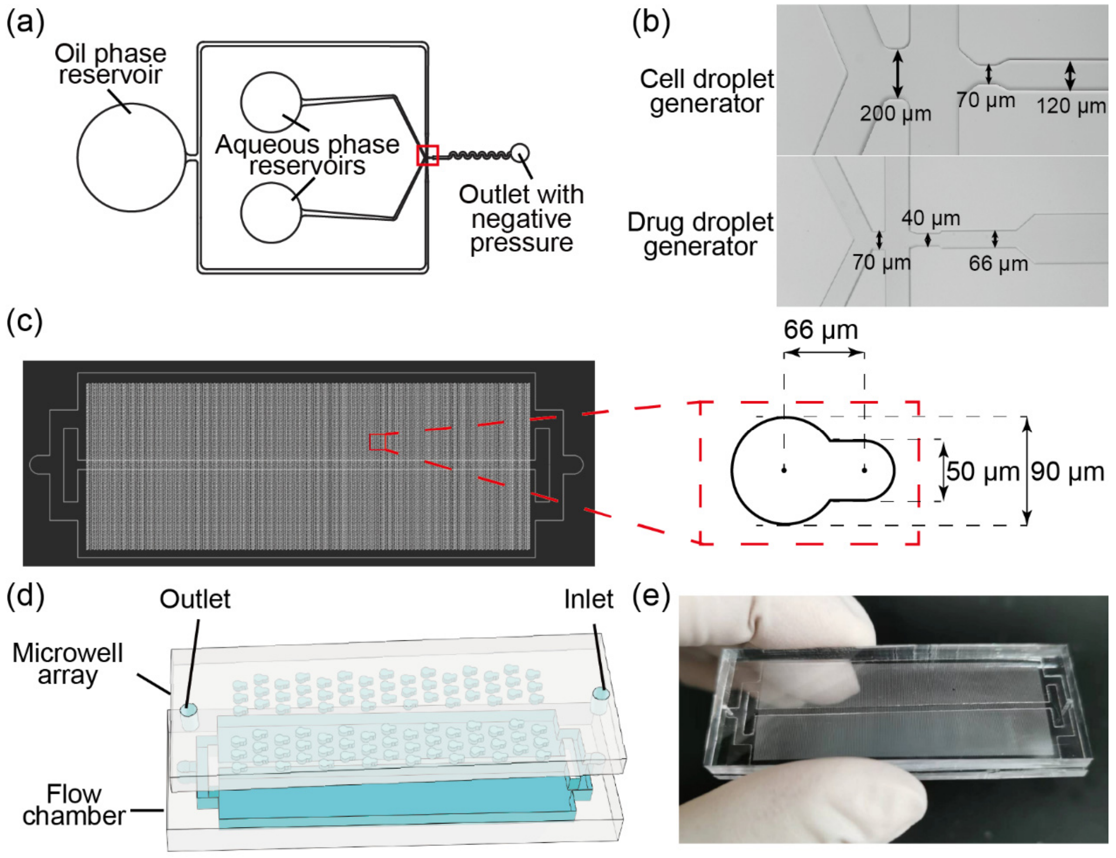


**Figure S1**. (a) Microfluidic chips of the droplet generators. (b) Close-up views with dimensions of the droplet generation region (boxed in red) in (a). The depths of the cell and drug droplet generators are 90 and 40 μm, respectively. (c) Design of the microwell array device and the dimension of the microwell unit. The depth of the microwell is 70 μm. (d) Schematic showing the microwell array device. (e) Photograph showing the microwell array device.

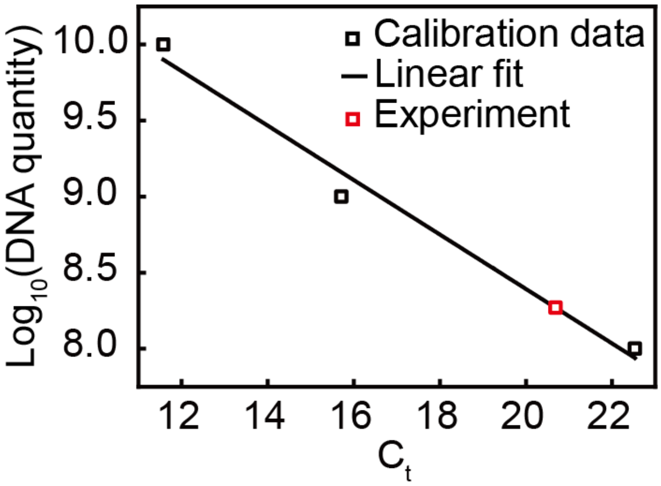


**Figure S2**. Quantity of the ssDNA in CADB complex recovered from 10,000 cells as a function of $C_t$ value in qPCR. Cell droplets were merged with droplets encapsulating a single type of CADB complex. After incubation, the cells are collected and lysed, and qPCR with primers (**Table S2**) targeting the ssDNA in the CADB complex were performed. By referring to the calibration curve (black line), the amount of ssDNA molecules immobilized on each cell was calculated to be 18621 ($10^{8.27}$ among 10000 cells), which was consistent with reported works.

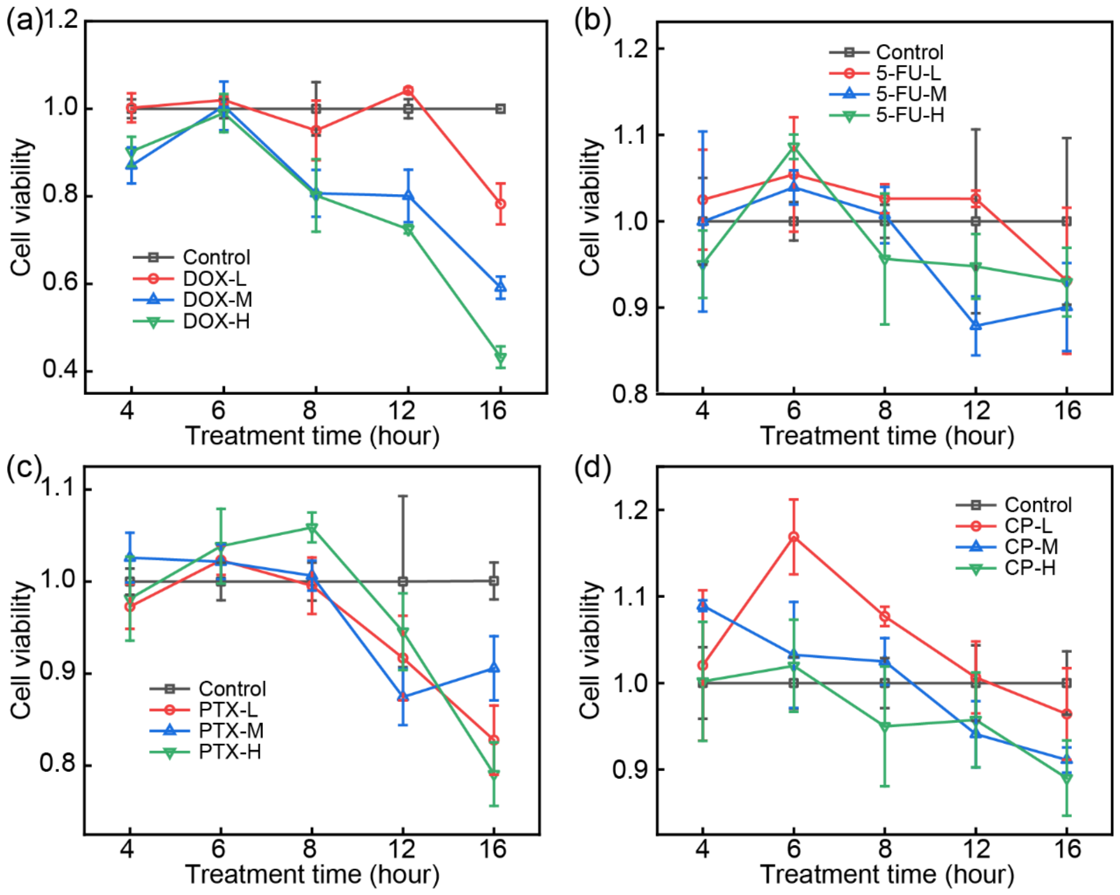


**Figure S3.** Cell viability tests using the CCK-8 assay to determine drug treatment time. MCF-7 cells were seeded in a 96-well plate ($1\times10^4$/well) with three replicates for each drug treatment condition.

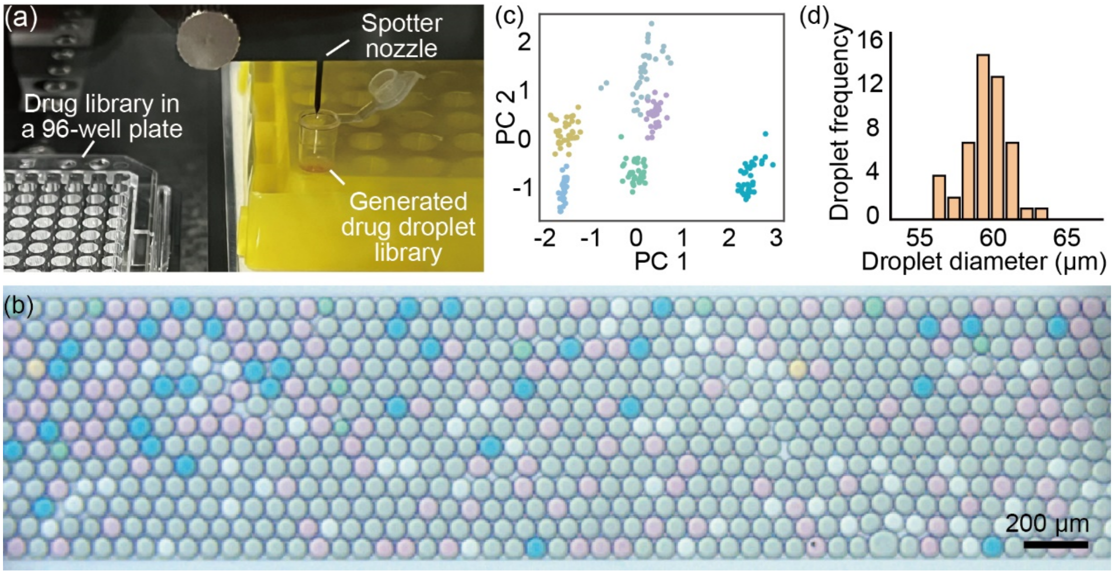


**Figure S4**. Generation of droplet libraries using a microarray spotter. (a) Photograph showing the key components of microarray spotter. (b) Micrograph showing the generated droplets. Six color-coded solutions were used. (c) Scatter plot of principal components of the RGB values showing that the six types of droplets can be recovered. (d) Diameter distribution of the generated droplets.

**Table S1.** Reagent, materials, and other consumable information.

| Reagent | Vendor | Catalog number |
|---|---|---|
| Dulbecco's modified Eagle's medium (DMEM) | Gibco | C11965500BT |
| Fetal bovine serum (FBS) | Gibco | 192-1005PJ |
| Penicillin-streptomycin (PS) | Gibco | 15140-122 |
| Trypsin-EDTA | Gibco | 25200056 |
| Ficoll-PM400 | Cytival | 17030010 |
| HEPES | Pluronic | 24040-032 |
| DPBS | Gibco | 14190-144 |
| Doxorubicin hydrochloride (DOX) | Sigma-Aldrich | D515 |
| Fluorouracil (5-FU) | Sigma-Aldrich | F6627 |
| Cyclophosphamide (CP) | Sigma-Aldrich | C0768 |
| Paclitaxel (PTX) | Aladdin | P106869 |
| Dimethylsulfoxide (DMSO) | Sigma-Aldrich | D8418 |
| Biotinylated concanavalin A | Sigma-Aldrich | C2272 |
| Recombinant streptavidin | BBI Solutions | C600432-0005 |
| AO/PI staining buffer | CountStar | N/A |
| Droplet generation oil | BIO-RAD | 1864005 |
| Polydimethylsiloxane (PDMS) | Dow Corning | 02085925 |
| 30 ml syringe | BD | 302833 |
| Cryovial | Corning | 431386 |
| 1H,1H,2H,2H–perfluoro-1-octanol (PFO) | Sigma-Aldrich | 370533 |
| Micro tubing | Scientific Commodities | BB31695-PE/2 |
| Qubit ssDNA HS assay kit | Invitrogen | Q10212 |
| Cell counting kit (CKK-8) | Yeasen Biotechnology | 40203ES76 |

**Table S2.** Concentrations of the drug groups (in µg/mL).

| Drug group | Concentration of the drug droplet | Concentration after merging |
|---|---|---|
| DOX-L | 68.5 | 10 |
| DOX-M | 342.3 | 50 |
| DOX-H | 684.6 | 100 |
| 5-FU-L | 8.9 | 1.3 |
| 5-FU-M | 44.5 | 6.5 |
| 5-FU-H | 178 | 26 |
| CP-L | 2740 | 400 |
| CP-M | 13692 | 2000 |
| CP-H | 27400 | 4000 |
| PTX-L | 34.2 | 5 |
| PTX-M | 68.4 | 10 |
| PTX-H | 136.8 | 20 |

**Table S3.** Design of the CADB barcode.

| **No.** | **Base sequences:** 5'-PCR handle-drug barcode-UMI-Poly(A)-3' | **Drug** |
|---|---|---|
| #1 | 5'-TTGTCTTCCTAAGACCGCTTGGCCTCCGACT-TAACAGCCAATCTGCGTAACAGCCAACCTTCC-NNNNNNNNNNB-AAAAAAAAAAAAAAAAAAAA | DOX-L |
| #2 | 5'-TTGTCTTCCTAAGACCGCTTGGCCTCCGACT-CTAAGAGTCCTCTGCGCTAAGAGTCCCCTTCC-NNNNNNNNNNB-AAAAAAAAAAAAAAAAAAAA | DOX-M |
| #3 | 5'-TTGTCTTCCTAAGACCGCTTGGCCTCCGACT-TTACTGCCTTTCTGCGTTACTGCCTTCCTTCC-NNNNNNNNNNB-AAAAAAAAAAAAAAAAAAAA | DOX-H |
| #4 | 5'-TTGTCTTCCTAAGACCGCTTGGCCTCCGACT-CGCTGAATTCTCTGCGCGCTGAATTCCCTTCC-NNNNNNNNNNB-AAAAAAAAAAAAAAAAAAAA | 5-FU-L |
| #5 | 5'-TTGTCTTCCTAAGACCGCTTGGCCTCCGACT-TGACGTCCTTTCTGCGTGACGTCCTTCCTTCC-NNNNNNNNNNB-AAAAAAAAAAAAAAAAAAAA | 5-FU-M |
| #6 | 5'-TTGTCTTCCTAAGACCGCTTGGCCTCCGACT-TGTGTGTAACTCTGCGTGTGTGTAACCCTTCC-NNNNNNNNNNB-AAAAAAAAAAAAAAAAAAAA | 5-FU-H |
| #7 | 5'-TTGTCTTCCTAAGACCGCTTGGCCTCCGACT-AGATTGAGAGTCTGCGAGATTGAGAGCCTTCC-NNNNNNNNNNB-AAAAAAAAAAAAAAAAAAAA | CP-L |
| #8 | 5'-TTGTCTTCCTAAGACCGCTTGGCCTCCGACT-GATAATGATGTCTGCGGATAATGATGCCTTCC-NNNNNNNNNNB-AAAAAAAAAAAAAAAAAAAA | CP-M |
| #9 | 5'-TTGTCTTCCTAAGACCGCTTGGCCTCCGACT-TTGAAGAAGCTCTGCGTTGAAGAAGCCCTTCC-NNNNNNNNNNB-AAAAAAAAAAAAAAAAAAAA | CP-H |
| #10 | 5'-TTGTCTTCCTAAGACCGCTTGGCCTCCGACT-CTGATACTTCTCTGCGCTGATACTTCCCTTCC-NNNNNNNNNNB-AAAAAAAAAAAAAAAAAAAA | PTX-L |
| #11 | 5'-TTGTCTTCCTAAGACCGCTTGGCCTCCGACT-GGCTCCAAGCTCTGCGGGCTCCAAGCCCTTCC-NNNNNNNNNNB-AAAAAAAAAAAAAAAAAAAA | PTX-M |
| #12 | 5'-TTGTCTTCCTAAGACCGCTTGGCCTCCGACT-AATCTACAAGTCTGCGAATCTACAAGCCTTCC-NNNNNNNNNNB-AAAAAAAAAAAAAAAAAAAA | PTX-H |
| #13 | 5'-TTGTCTTCCTAAGACCGCTTGGCCTCCGACT-TGAGGTGGTTTCTGCGTGAGGTGGTTCCTTCC-NNNNNNNNNNB-AAAAAAAAAAAAAAAAAAAA | Control |

**Table S4.** Oligo sequence and primers for the qPCR validation of the CADB binding and recovering.

| **Name** | **Base sequences (5' to 3')** |
|---|---|
| Oligo | 5'-GCTGCGCTCGATGCAAAATAGGTTTACTBAAAAAAAAAAAAAAAAAAAAAAAAAAAAAA |
| RT-primer | 5'-CACGCACTGACTGACAGACTTTTTTTTTTTTTTTTTTTTTTTTTTTTTTV |
| FP | 5'-GCTGCGCTCGATGCAAAATA |
| RP | 5'-CACGCACTGACTGACAGACT |

**Video S1**. A video showing the operation of droplet capturing, pairing, merging, and retrieval.
